# Priors for natural image statistics inform confidence in perceptual decisions

**DOI:** 10.1101/2023.02.19.529102

**Authors:** Rebecca K West, Emily J A-Izzeddin, David K Sewell, William J Harrison

## Abstract

Decision confidence plays a critical role in humans’ ability to make adaptive decisions in a noisy perceptual world. Despite its importance, there is currently little consensus about the computations underlying confidence judgements in perceptual decisions. In order to better understand these mechanisms, in this study we sought to address the extent to which confidence is informed by a naturalistic prior probability distribution. Contrary to previous research, we did not require participants to internalise the parameters of an arbitrary prior distribution. Instead we used a novel psychophysical paradigm which allowed us to capitalise on probability distributions of low-level image features in natural scenes, which are well-known to influence perception. Participants reported the subjective upright of naturalistic image target patches, and then reported their confidence in their orientation responses. We used computational modelling to relate the statistics of the low-level features in the targets to the distribution of these features across many natural images. As expected, we found that participants used an internalised prior of the regularities of low-level natural image statistics to inform their perceptual judgements. Critically, we also show that the same low-level image statistics predict participants’ confidence judgements. Overall, our study highlights the importance of using naturalistic task designs that capitalise on existing, long-term priors to further our understanding of the computational basis of confidence.

## Introduction

Humans make countless decisions based on noisy perceptual information every day, and these decisions are often accompanied by a sense of confidence. This sense of confidence, defined as the belief that a choice or proposition is correct based on the available evidence (Pouget et al., 2016), plays a crucial role in humans’ ability to make adaptive decisions. For example, confidence has been shown to influence subsequent decisions (van den Berg, Zylberberg, et al., 2016) and to guide the search for further information before committing to a choice (Desender et al., 2018). Despite being a salient property of decision making, there is currently little consensus on how humans compute their confidence. To better understand the computations involved in generating decision confidence, the aim of the current study was to investigate the extent to which confidence is informed by *prior knowledge*. Specifically, we investigated how *long-term* priors for the statistical distribution of low-level visual features, formed either explicitly or implicitly through experience with the natural world, influences confidence.

Several models have been proposed to describe how humans compute their confidence in their perceptual decisions. One prominent theory suggests that confidence is computed according to the rules of Bayesian inference (Adler & Ma, 2018; Aitchison et al., 2015; Bertana et al., 2021; Hangya et al., 2016; Li & Ma, 2020; Lisi et al., 2021; Locke et al., 2022; Navajas et al., 2017; Sanders et al., 2016). According to Bayesian decision theory, observers combine prior beliefs about the state of the world (the *prior*) with the present sensory input (the *likelihood*) to compute a posterior probability distribution (Drugowitsch et al., 2014b; Hangya et al., 2016; Kepecs and Mainen, 2012; Meyniel et al., 2015; Pouget et al., 2016). The posterior probability distribution, thus, represents all possible observations and their inferred probabilities. To form a single representation or choice, an ideal Bayesian observer selects the most likely observation from the posterior probability distribution. Confidence can then also be derived from this posterior probability distribution and can be defined as the degree of certainty (or probability) associated with the representation that the observer has chosen (or intends to choose; Ma & Jazayeri, 2014).

In contrast to Bayesian models, several alternative models of confidence have been proposed. These models posit that confidence reports are better explained by suboptimal or *heuristic* computations, suggesting that confidence is based on learned associations between certain stimulus cues and specific outcomes. These models have received some empirical support with a range of studies linking confidence to heuristics such as external noise (Bertana et al., 2021; Boldt et al., 2017; Spence et al., 2016), response time (Faivre et al., 2018; Patel et al., 2012; van den Berg, Anandalingam, et al., 2016), stimulus familiarity (Reinitz et al., 2012) and task difficulty cues (Mole et al., 2018).

Despite substantial research interest and myriad explanatory models, the computations underlying the generation of decision confidence remain unclear. Empirical studies investigating the plausibility of Bayesian and non-Bayesian models of confidence have yielded mixed results, with some studies finding support for Bayesian models (Aitchison et al., 2015; Li & Ma, 2020; Navajas et al., 2017; Sanders et al., 2016) and others for non-Bayesian models (Adler & Ma, 2018; Aitchison et al., 2015; Bertana et al., 2021; Denison et al., 2018; Lisi et al., 2021; Locke et al., 2022; West et al., 2023). Furthermore, one of the major limitations of existing research is that many previous studies have exposed participants to stimuli drawn from arbitrarily pre-specified prior distributions, and compared their behaviour to that of a Bayesian optimal observer with full knowledge of that distribution (Denison et al., 2018; Li & Ma, 2020; Locke et al., 2022; Qamar et al., 2013; West et al., 2023). Such an approach is limited by the ability of humans to internalise the statistics of the prior distribution within the time frame of the experiment (Girshick et al., 2011). In order to better understand the computations underlying confidence judgements, we therefore sought to investigate the extent to which confidence is informed by a prior probability distribution that is derived from the statistics of natural environments, a *naturalistic prior.* Following a recent study (A-Izzeddin et al., 2024), we took advantage of the distributions of low-level features present in naturalistic stimuli to define a prior distribution, the statistics of which have been shown to bias humans’ perceptual decisions (Appelle, 1972; Berkley et al., 1975; Dakin, 2001; de Gardelle et al., 2010; Girshick et al., 2011).

A-Izzeddin and colleagues (2024) introduced a novel psychophysical paradigm which shows a clear link between environmental statistics and perceptual inferences. A-Izzeddin and colleagues (2024) presented participants with randomly oriented target images of outdoor scenes and participants were required to infer their subjective “upright” orientation. The targets were designed so that they were windowed within a small aperture of the original image, meaning that the targets themselves contained very little high-level structure. Participants, therefore, had to rely on a strategy which depended on the low-level image features in the targets to judge “upright”. The authors sought to determine if participants had an internal representation of the distribution of low-level features in natural images, a prior for environmental statistics, which they could use to guide their judgements. For example, as shown in **Figure 1**, horizontal orientation features, and, to a lesser extent, vertical orientation features are over-represented in natural images.

**Figure 1.**
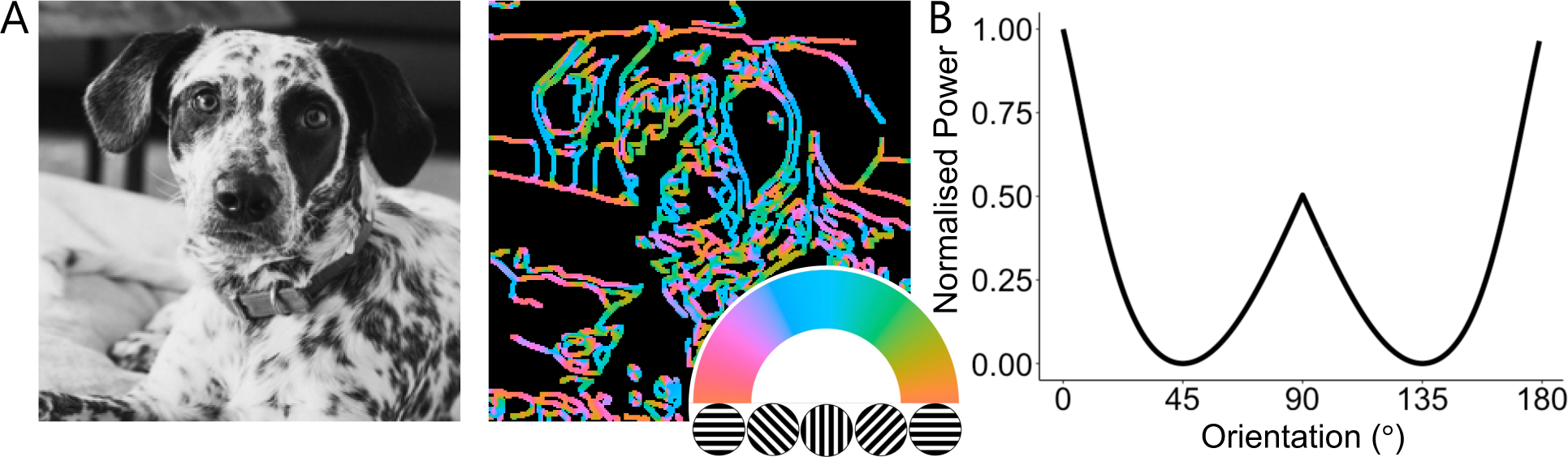
Natural image statistics. **(A)** Orientations of edges in digital photos have systematic biases. Original photo taken by Rafael Forseck and used under the Unsplash Licence. **(B)** Idealised distribution of orientations across many natural images (Hansen & Essock, 2004; Harrison et al., 2023).

Consistent with an internalised prior for environmental statistics, A-Izzeddin and colleagues found that participants’ distribution of orientation responses was well approximated by the frequency distributions of orientations in natural images. The authors used a model observer to show that participants’ responses were best explained by a process in which they oriented the targets so that the low-level features in the targets, such as orientation and phase, best approximated the *prior* distribution for these features in the environment. This study, therefore, is similar to others that demonstrate that humans have an *existing, internal model of a prior probability distribution of low-level image statistics* which they use to inform their perceptual inferences about the world (A-Izzeddin et al., 2024; see also Girshick et al., 2011). Hence, in the present study, we used the same paradigm to investigate confidence because it allowed us to determine the extent to which participants use an internal prior distribution to compute their confidence without requiring them to explicitly learn the parameters of that distribution.

In the present study, participants performed a perceptual task in which they made an orientation judgement about a target image and then reported their level of confidence in that orientation response (see **Figure 2A**). To investigate whether participants’ perceptual and confidence judgements were consistent with an internalised representation of natural image statistics, we used image processing techniques in combination with computational modelling. To preview our results, we confirmed that participants adopted a strategy that aligned the distribution of oriented features and phase with a prior probability model of the distribution of those features in the natural world (A-Izzeddin et al., 2024). Importantly, we also found that participants used the same prior probability distribution to inform their confidence judgements, *even without explicit instruction about the prior*. Our findings highlight the importance of naturalistic, long-term priors for the computation of decision confidence.

**Figure 2.**
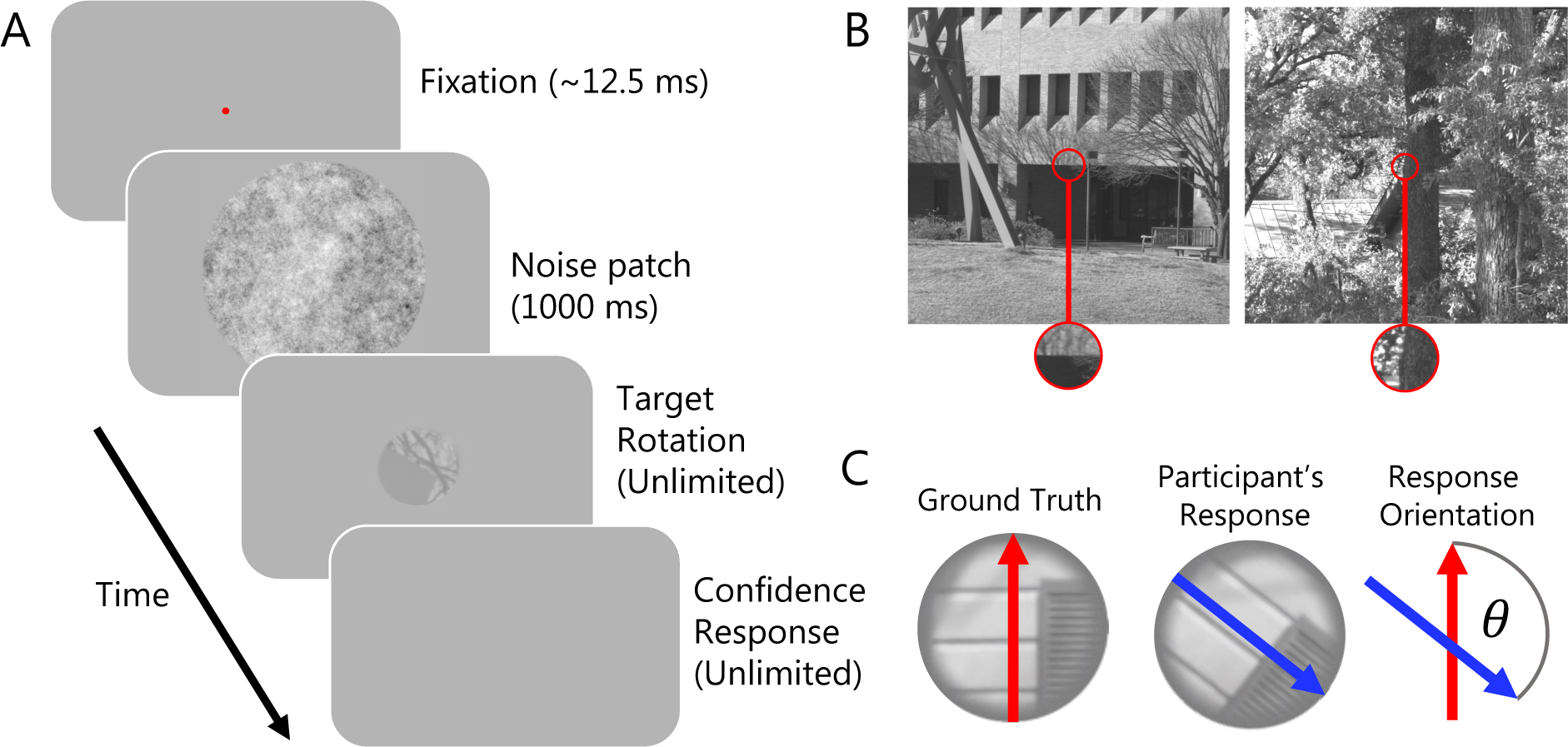
Experimental paradigm. **(A)** Schematic showing trial structure. Participants saw a fixation point, followed by a noise patch and then a target was presented at a random orientation. Participants used the mouse to rotate the target to appear upright and then made a confidence judgement (either “low confidence” or “high confidence”) in their chosen orientation response. **(B)** Targets, such as the examples outlined in red, were extracted from high quality photos of natural scenes (Burge & Geisler, 2011). **(C)** To quantify perceptual performance, we computed the angular difference between the objective upright orientation of the target and the participants’ chosen orientation.

## Methods

### Overview

On each trial, a participant viewed a randomly oriented *target*, positioned in the centre of the display (see **Figure 2A**). The participant was informed that the target was a circular patch that had been cropped from the centre of a larger *source image,* where source images were randomly selected from a database of images of outdoor scenes (see **Figure 2B**). The participant was instructed to rotate the target to the “upright” orientation, with no additional contextual information given about the source image. For each target, the participant made their orientation judgement and then made a confidence judgement in their chosen orientation response, reporting either “high confidence” or “low confidence”.

### Participants

In total, 21 participants (*M_age_* = 23.95, *SD_age_* = 4.65; no exclusions) completed the experiment. 10 participants completed Study 1 and 11 participants completed Study 2. All methods were the same for Study 1 and Study 2, unless indicated otherwise. As there did not appear to be any clear differences in the results of Study 1 and Study 2, we combined datasets across studies. We, therefore, report results for all 21 participants. See **Supplementary Material: Experiment 1 and Experiment 2 Comparison** for additional commentary. All participants were naïve to the purpose of the study and were reimbursed for their time ($20 per hour in cash). Ethics approval was granted by the University of Queensland Medicine, Low & Negligible Risk Ethics Sub-Committee.

### Stimuli & Apparatus

All participants saw the same 500 digital natural images, selected randomly from a database of high-resolution photos of outdoor scenes and were cropped to 1080×1080 pixel regions (Geisler & Perry, 2011). Targets were circular patches cropped from the centre of the 1080×1080 images, subtending 2° of visual angle in diameter (see **Figure 2B**). All stimuli were converted to greyscale using MATLAB’s rgb2gray() function. During practice, targets were selected randomly from a different set of images from the same database.

Stimuli were presented using the Psychophysics Toolbox (3.0.12; Brainard, 1997; Pelli, 1997) and a gamma correction was applied to the display, assuming gamma was 2. In Study 1, stimuli were presented on a 24-inch ASUS VG428QE monitor, 1920 x 1080-pixel resolution and a refresh rate of 100 Hz. Participants were seated in a dark room with their head positioned on a chin rest fixed at a viewing distance of 57cm. In Study 2, stimuli were presented on a 24-inch DELL P2414H monitor, 1920 x 1080-pixel resolution and a refresh rate of 60 Hz.

### Procedure

As shown in **Figure 2A**, at the start of each trial participants were presented with a central fixation point for ∼125 ms (fixation time sampled from a uniform distribution between 0 and 250 ms) followed by a black and white pink noise patch for 1000 ms (27° of visual angle in diameter). The target was then presented centrally at a random orientation and participants used the mouse to re-orient the patch to be “upright”. Participants clicked the mouse to confirm their response and then used the arrow keys (left arrow = low confidence and right arrow = high confidence) to indicate their confidence in their chosen orientation. Half of the participants were instructed to use the high confidence and low confidence ratings approximately equally often (Study 1) and the other half of participants received no additional instructions about using the confidence ratings (Study 2). All participants completed 500 trials with the same targets, the order of which was randomised for each participant. Trials were split into 5 blocks of 100 trials with self-paced breaks in between blocks.

Prior to completing the experiment, participants did 40 practice trials to familiarise themselves with the task. In the first 20 practice trials, participants only rotated the targets to their “upright” orientation and, in the remaining 20 trials, participants rotated the targets and then made confidence ratings. Participants saw different target images during testing and training.

### Data Analyses

We used statistical models and digital signal processing techniques (e.g. Harrison, 2022) to investigate whether participants use a prior for natural image statistics when making perceptual inferences. We then developed a novel statistical model to determine if participants use the same prior to inform their confidence judgments. In the following sections, we first describe the image processing methods used to derive the distribution of orientation energy and phase for the prior and each target patch. We then describe the perceptual and confidence models.

The targets were designed so that they were windowed within a small aperture of the larger source image. This meant that the targets themselves contained very little high-level structure that participants could use to unambiguously infer the upright orientation of the target (see **Supplementary Material: High-Level Image Features and “Informativeness” Data** for further discussion on this issue). Participants, therefore, had to rely on a strategy that depended on the low-level image features in the targets only (see **Supplementary Figure 8**). Based on A-Izzeddin and colleagues (2024), we expected that participants would adopt a strategy in which they chose rotational offsets for the targets which best matched the distribution of low-level features in the targets to the distribution of these features in the environment, referred to as the *prior*. Consistent with A-Izzeddin and colleagues (2024), we focused on the use of two specific low-level features, the distribution of orientation energy and phase, and defined the prior as the distribution of these features across thousands of images of natural scenes (see **Figure 1**). Importantly, for the experimental task, we did not provide any guidance on what features participants should use to make their decisions or give them any explicit instructions about the prior. Instead, we relied on participants’ *internal* representation of the prior to do the task.

#### Orientation Energy

##### Prior

We defined the prior distribution of orientation energy based on studies of natural images (e.g. Wei & Stocker, 2015):

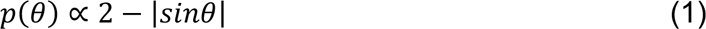

where *p*(*θ*) is the probability of observing orientation energy with an orientation of σ in radians. **Equation 1** assumes equal prevalence of horizontal and vertical orientations. Other studies, however, have shown that horizontal features are over-represented relative to vertical features (Hansen et al., 2003; Hansen & Essock, 2004; Harrison, 2022). We therefore modified **Equation 1** by increasing the proportion of horizontal energy to approximately match the asymmetry of vertical and horizontal features in natural images. We did this by multiplying the prior shown in **Equation 1** by a von Mises function so that:

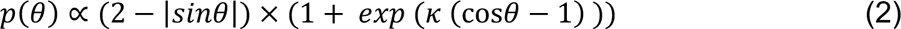

where *κ* is the width of the von Mises function which we set to 2.5. *p*(*θ*) was then normalised within the range 0 – 1 so that it matched the range of normalised energy in the targets. Small changes in *κ* did not change the results. **Figure 1** shows the prior distribution of orientation energy (black distribution).

##### Target Patch

We calculated the distribution of orientation energy in the target to compare against the prior. For a given target patch, we computed orientation energy in 180 equally space orientation bands, each of which covered all spatial frequencies. These operations were performed in the frequency domain; energy was the absolute of the Fourier-transformed target patches. Orientation filters were also constructed in the frequency domain as raised cosine filters with a bandwidth of 45° (full width half height). Energy was summed within each orientation band, giving a distribution of energy across orientations. Because absolute energy fluctuates from one image to the next, the distribution of energy was normalised within the range 0 – 1, by subtracting the minimum value, and then dividing by the maximum value. **Figure 4B/D** shows example distributions of orientation energy (orange distribution) for an example target patch at two different rotational offsets in **Figure 4A/C**.

#### Phase

As described in A-Izzeddin et al. (2024), contrast energy alone is circular around ±90°, whereas observers’ reports are circular around ±180°. To estimate fully circular responses, therefore, a second image statistic is required to determine whether an image needs to be inverted. We modelled this process as an estimate of the lighting direction in the target using the response of a phase-locked filter.

##### Prior

Consistent with previous research, we assumed that participants had a light-from-above prior (Adams, 2007; Brewster, 1826; Metzger, 1936; Murray, 2013; Ramachandran, 1988). We operationalise lighting direction as the image’s phase distribution (i.e., the response of a phase-locked filter at all orientations). To instantiate the light-from-above prior computationally, we quantified the distribution of phase-locked filter responses using the entire stimulus set. We convolved a bank of sine wave filters, whose orientations ranged from 0° - 179°, with each target patch, and then found the average of the filter responses within the centre of the target patch to exclude phase artifacts introduced by the circular aperture. The resulting mean phase distribution was clearly sinusoidal (**Supplementary Figure 10**). We therefore modelled this phase distribution using the best fitting sinusoid. This sinusoid was defined by the following equation:

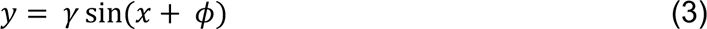

Where *γ* is a scaling factor and *ϕ* is an offset. We fit the model using ordinary least squares, capturing 97% of variance in mean filter responses. The best fitting value for *ϕ* was 41° (i.e., relative to a normal sinewave, the model was phase-shifted by 41°). We therefore treat the function *y* = sin(*x* + 41) as the prior distribution of lighting as detected by phase filters.

##### Target Patch

We also calculated the phase distribution in the target to compare against the prior distribution. The phase distribution was estimated for each target by convolving phase-locked sine wave filters at each orientation. **Figure 4** shows example distributions of phase.

### Perceptual Model

We used the model observer developed by A-Izzeddin and colleagues (2024) to investigate whether participants use an internal prior model for low-level image features to inform their orientation judgements. This model is defined as a “pretty good observer” model because it uses a subset of image statistics to make a decision, as opposed to an “ideal observer” model which exploits all possible sources of information.

The model included two stages. In the first stage, we rotated the target to best match the distribution of orientation energy in the target with the prior. We used MATLAB’s fminsearch() function to find the rotational offset that minimised the sum of squared differences in each orientation band between the target and the prior (see **Figure 4**). To avoid local minima, we fit the model with varying starting parameters and used the rotational offset at the global minimum from all fits. Because orientation contrast is phase invariant, in the second stage of the model after finding the best rotational offset, we estimated the phase distribution in the target using phase-locked filters and inverted the target so that it provided the best match to the prior distribution of phase in natural images. That is, so that the lighting direction in the target was consistent with a light-from-above prior. To determine the correct inversion, the prior distribution of phase was regressed onto the phase distribution in each target patch to estimate a scaling factor *γ*. A negative *γ* value indicated that the target needed to be inverted to best match the aggregate prior phase distribution and hence, led the model observer to invert the target relative to the orientation identified in the orientation prior stage of the model. For non-negative *γ* values, the model observer left the target at the orientation identified in the orientation prior stage of the model.

The fitting of the perceptual model as described thus far was independent from participants’ responses – we fit the targets’ statistics to the prior. This meant that the model was deterministic. Because all participants saw the same targets, the model predicted the same pattern of responses for all participants. To account for deviations from the model observer across participants, we fit a single noise parameter for each participant: we added a random amount of noise, *ε*, to the model’s predicted response orientation for each target. The amount of added noise was sampled from a normal distribution with a mean of 0 and a standard deviation, *σ*, that was estimated separately for each participant, *j*, such that:

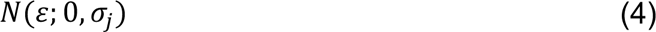

where *N* denotes the normal density function and *ε* is the amount of noise added to the perceptual model’s prediction. We estimated *σ* by minimising the difference between each participant’s observed distribution of response orientations and the mean predicted distribution of response orientations for that participant using ordinary least squares according to:

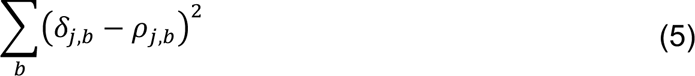

Where *δ* is the observed proportion of responses and *ρ* is the predicted proportion of responses in bin, *b*, for participant, *j*. To calculate the predicted proportion of responses, we simulated 1000 noisy responses by drawing random samples of *ε* according to **Equation 4** for each trial and each participant and added these random samples to the model’s predicted orientation response for that target. We then calculated the average proportion of responses in each orientation bin for each participant. See **Supplementary Table 4** for *σ* parameter estimates.

### Confidence Model

For the associated confidence judgements, we sought to determine if participants use the same prior for the distribution of low-level features in natural scenes to inform their confidence. If participants use the same prior to inform their confidence, we hypothesised that their confidence judgements would reflect how closely the distribution of low-level features in their chosen orientation response matched the prior distribution. To test this prediction, we developed several measures that summarised the degree of overlap in orientation energy and phase in the target, as oriented by the participant, and the prior (described above). In contrast to these prior-related measures, we also considered the possibility that confidence depends on other stimulus cues not related to the prior, such as the contrast of the target patch or the response time of the perceptual orientation judgement. To evaluate these predictions, we used a generalised linear mixed model framework to investigate the relationship between certain features of the target and participants’ confidence judgements. Importantly, we computed the measures (described in detail below) using the targets at the rotational offset chosen by the participant on each trial, reflecting the fact that confidence judgements are a second-order reflection on the first-order decision.

In the sections below, we first describe the general modelling framework (the generative model) and then describe each of the measures used to predict confidence.

#### Generative Model

We used a generalised linear mixed model (GLMM) in the form of a modified logistic regression to relate a set of weighted predictors to binary confidence responses (low or high). In this framework, the sum of weighted predictors is passed through a logistic link function which transforms the unbounded weighted sum into the range of [0, 1]. The linear function is given by:

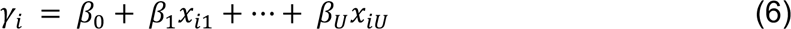

where *x_i_*_1_ … *x_iU_* are the values for the *U* predictors on trial *i, β*_1_ … *β_U_* are the weights for the respective predictors and *β*_0_ is the intercept term. The inverse logistic link function that the linear function is passed through is given by:

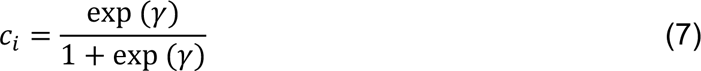

We used a multilevel extension of this model, such that we estimated the predictor weights, *β*s, for each participant. These weights are assumed to come from the same population such that the individual-participant *β*s are estimated concurrently with the mean and variance of the weights at the population level (Wallis et al., 2015). For the confidence model, we used a model with 8 predictors, where the linear function was described by:

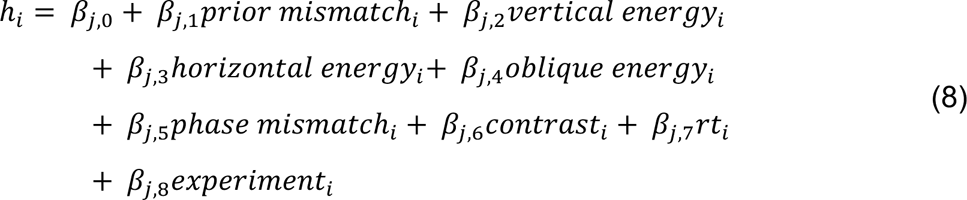

such that for participant *j*, *β_j_*_,1_ … *β_j_*_,5_ were the weights for the respective predictors and *β_j_*_,0_ was the intercept. *β_j_*_,8_ was the weight for an experiment indicator variable which allowed for differences in mean confidence across experiments (see **Supplementary Figure 1**). All predictors were standardised, by subtracting the mean and dividing by the standard deviation, prior to model fitting (see **Supplementary Figure 7**). The computation of each predictor is described below.

#### Orientation Energy

The goal of the confidence model was to determine if participants’ confidence judgements were informed by the same prior used for their perceptual judgements. We reasoned that if participants used the same prior to inform their confidence, there would be a statistical association between confidence and the amount of correspondence in the distribution of low-level features in the target and the prior. To summarise the degree of overlap between the distribution of orientation features in the target and the prior, we used two sets of predictors: *orientation*-*prior-mismatch* and *cardinal and oblique orientation energy*. For both sets of predictors, we assumed that participants use prior knowledge about the statistics of the distribution of orientation features in natural scenes to inform their confidence. For the orientation-prior-mismatch predictor, we assumed that observers directly compare a veridical representation of the distribution of orientation energy in the stimulus and veridical knowledge of the prior distribution of orientation energy to compute their confidence. This means that observers use a statistic that summarises the mismatch between the *entire* distribution of orientation energy in the target and the prior to compute their confidence. For the cardinal and oblique orientation energy predictors, in contrast, we assume that observers rely on only a *subset* of features in the stimulus and the prior model to compute their confidence. Specifically, observers use a set of salient orientation cues, namely vertical, horizontal, and oblique features, for confidence, based on prior knowledge about the probability of these orientation features in the natural environment.

##### Orientation-Prior-Mismatch

For the orientation-prior-mismatch predictor, we calculated the average difference between the distribution of orientation energy in the target at the rotational offset chosen by the participant and the prior distribution of orientation energy. We did this by calculating the difference in orientation energy between the target and the prior in each orientation bin, squaring this difference and then summing across bins.

##### Cardinal and Oblique Orientation Energy

For the cardinal and oblique orientation energy predictors, we assumed that participants use a subset of orientation features to estimate their confidence. We calculated orientation energy across all orientations for each target positioned at the rotational offset chosen by the participant. We then normalised energy for each orientation band so that each band expressed a proportion of orientation energy in that bin relative to total orientation energy in the target, a form of divisive normalisation (Carandini & Heeger, 2012; see **Supplementary Figure 6**). To predict confidence, we use 3 predictors: *vertical energy* (where 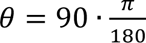), *horizontal energy* (where *θ* = 0 *or* 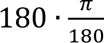), and *oblique energy* (where 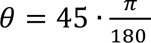 *or* 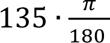).

#### Phase

To summarise the degree of overlap between the distribution of phase in the target and the prior, we used a single predictor: *phase*-*prior-mismatch*.

##### Phase-Prior-Mismatch

For the phase-prior-mismatch predictor, we computed the difference between the scaled phase distribution in the target and the prior distribution of phase at each filter orientation (0° - 179°, see **Equation 3**). We squared the difference for each filter orientation and then summed across orientations to have a single measure which summarised the difference between the phase distribution in the target and the prior.

#### Response Time

As described above, we also considered the possibility that confidence depended on other heuristic cues not related to the prior. Based on previous research (Faivre et al., 2018; Patel et al., 2012; van den Berg, Anandalingam, et al., 2016), we therefore included a predictor in the confidence model for *response time*. For the response time predictor, we used the response time of the orientation response for each trial, measured in milliseconds.

#### Contrast

We also postulated that participants may use the contrast of the target as a heuristic cue for confidence. For the *contrast* predictor, we used the root-mean-square (RMS) contrast of each target (Bex & Makous, 2002; Harrison, 2022). RMS contrast is the standard deviation of the luminance (i.e., pixel) values:

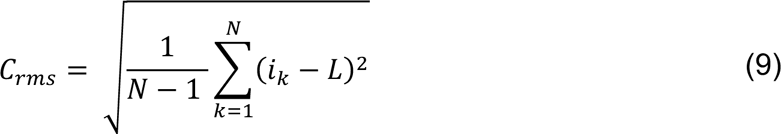

where *k* is a pixel index, *N* is the total number of pixels, and *L* is the mean luminance.

#### Summary of Modelling Approach

To summarise our expectations about the confidence modelling results, we reasoned that if participants used a natural image prior to inform their confidence, we would find that the measures which summarise the degree of overlap in orientation energy and phase between the target and the prior (orientation-prior-mismatch and/or cardinal/oblique orientation energy and phase-prior-mismatch) predicted confidence. If, participants used other non-prior related measures to inform their confidence, either in addition to or instead of the prior related measures, we would expect the other heuristic cues, such as response time or contrast, to predict confidence.

## Results

To better understand the computations underlying decision confidence, we investigated whether confidence judgements in perceptual decisions are informed by priors. Rather than requiring participants to learn the statistics of an arbitrary, contextual prior distribution, we used a long-term, naturalistic prior distribution that has been previously shown to affect participants’ perceptual inferences: the distribution of orientation energy and phase in natural scenes. To validate this approach, in our first analysis, we confirmed a strong influence of the prior on participants’ perceptual judgements as recently reported (A-Izzeddin et al., 2024).

### Perceptual Judgements

To quantify perceptual performance, we computed the angular difference between the objective upright orientation of the target and the participants’ chosen orientation, referred to as the *response orientation* (see **Figure 2C**). Response orientation was measured in degrees ranging from −180° to 180°. The black line in **Figure 3A** shows the frequency distribution of response orientations. If participants (*N* = 21) were not able to infer the objective upright orientation of the image using the low-level features in the target, the distribution of response orientations would be distributed uniformly across the range −180° to 180°. By contrast, **Figure 3A** shows that the most frequent response orientation was 0°, suggesting the modal response was highly accurate, and there are clear peaks at the cardinal orientations (±90° and ±180°), where targets were either inverted relative to their true orientation or offset by 90°. The presence of peaks aligned to cardinal orientations in **Figure 3A** is generally consistent with observers aligning edges within the target to the most common orientations found in nature. Note, however, that observers made ±180° inversions less frequently than 0° reports, which requires more than simply aligning the target energy to an orientation prior (in which case there would be an equal proportion of 0° and ±180° responses). To further investigate how participants made their responses, we used a model observer, described below.

**Figure 3.**
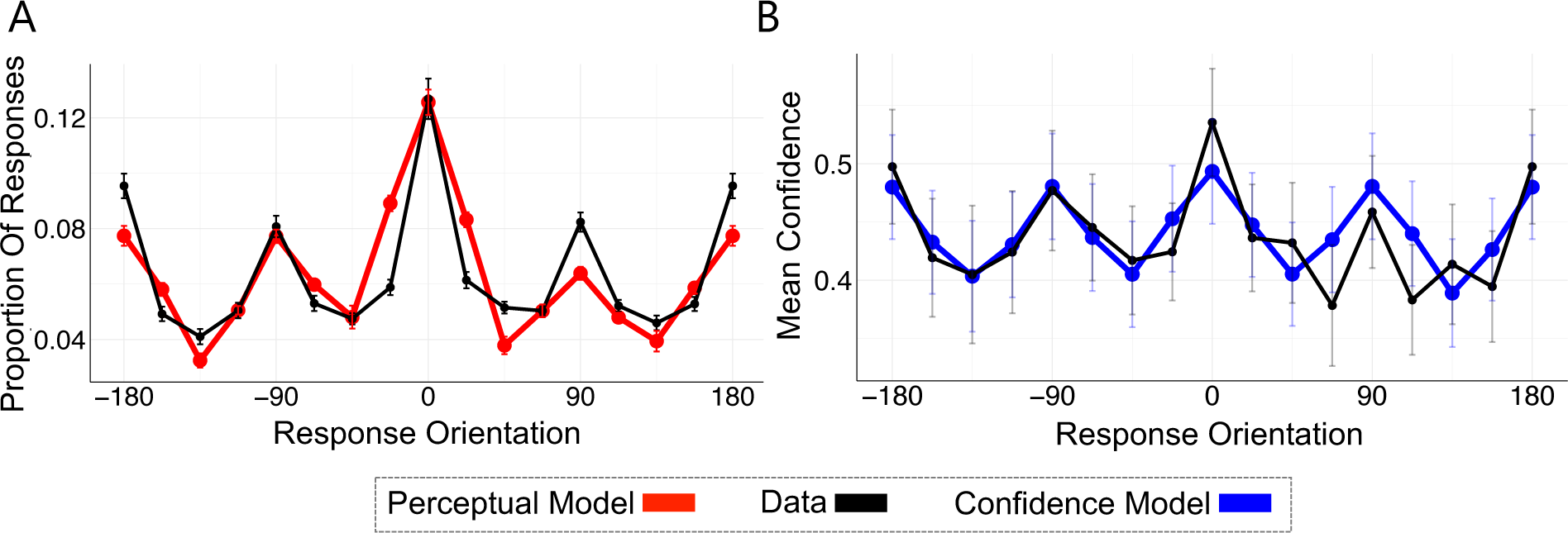
Perceptual and Confidence Data and Models. **(A)** The distribution of observers’ binned response orientations is shown in black. The output of the perceptual model, averaged across observers, is shown in red, and provides a good fit to the empirical data. **(B)** Mean confidence across binned response orientations shown in black. The output of the confidence model is shown in blue, and provides a good fit to the empirical data. Bin size = 22.5°. Error bars show ± 1 *SEM*.

#### Perceptual Model

We modelled participants’ orientation judgements with a “pretty good observer model” developed by A-Izzeddin and colleagues (2024; see **Methods** for detailed model description). Whereas an *ideal* observer model exploits all possible sources of information to make a decision, the pretty good observer model was designed to use only a subset of image statistics to make a decision. The model included two stages. In the first stage, the model rotated the target to best match the distribution of orientation energy in the target to the prior distribution (see **Figure 4**). In the second stage, the model used the target’s phase distribution to estimate lighting direction. As described in more detail by A-Izzeddin et al., the motivation of this second stage was to approximate a light-from-above prior (Adams, 2007; Brewster, 1826; Metzger, 1936; Murray, 2013; Ramachandran, 1988). This two-stage procedure produced a pattern of modelled response orientations for the 500 targets shown to each participant. To account for deviations from the model observer across participants, we fit a single noise parameter for each participant (see **Methods – Perceptual Model**).

**Figure 4.**
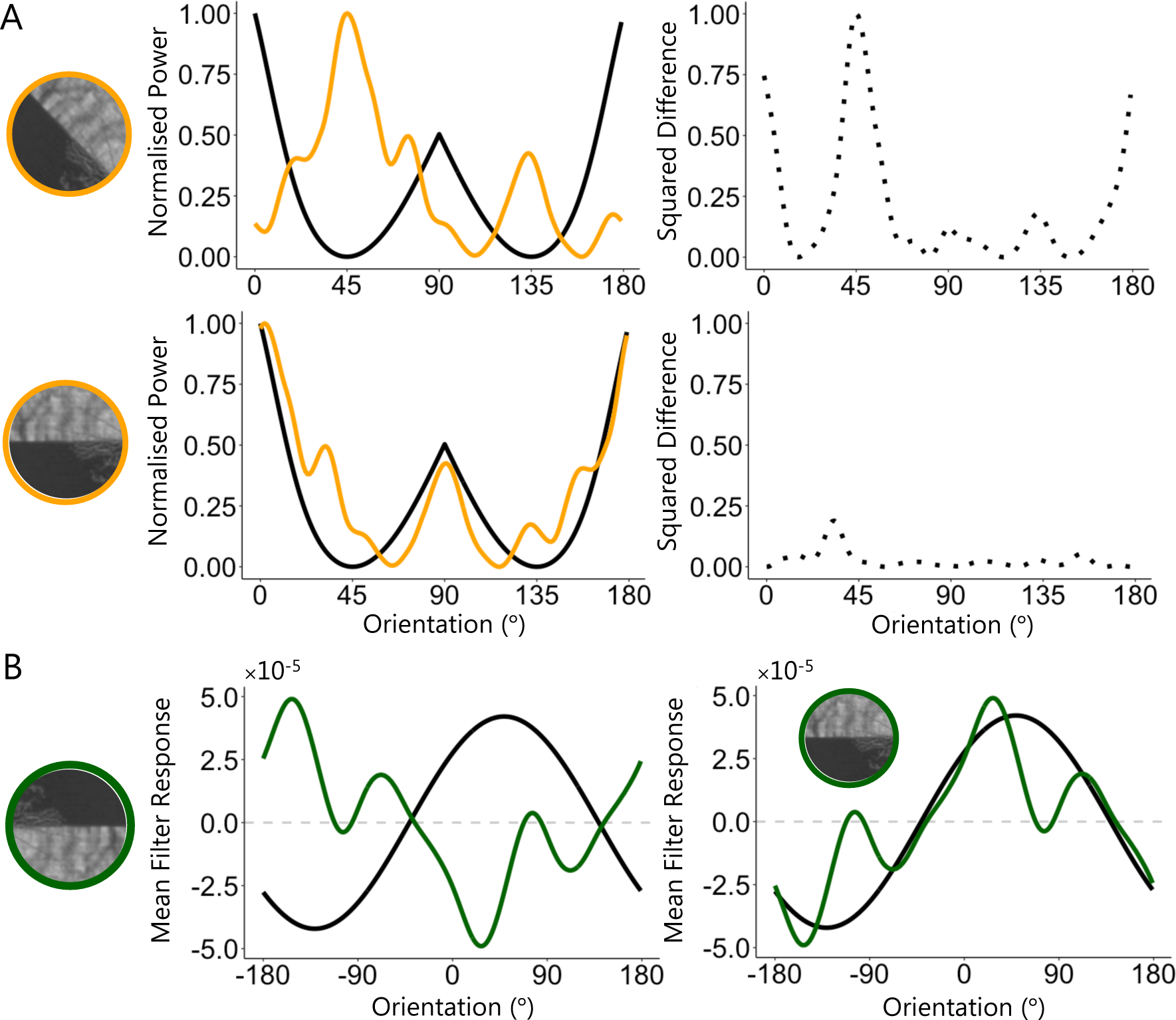
First and Second Stage of the Perceptual Model. Left discs show an example target patch at different orientations, while the right panels show the corresponding orientation energy (orange) and phase distributions (green) compared with the prior (black). **(A)** In the first stage of the perceptual model, the target is oriented to minimise the sum of squared differences between the distribution of orientation energy in the target and the prior. Right panels show the squared difference in normalised power (dotted black distribution) between the target and the prior for each orientation bin. **(B)** In the second stage of the perceptual model, the distribution of phase in the target is compared to the phase prior. Green distributions show phase distribution for an upright (left) and inverted (right) target compared with the prior (solid black distribution; i.e., the light-from-above prior).

As shown in **Figure 3A**, the perceptual model provided a very good approximation of participants’ responses. The close correspondence between participants’ responses and model responses suggests that participants judge the orientation of the target by matching a relatively simple set of low-level image statistics, namely orientation energy and phase, to an internal representation of the distribution of these statistics in the natural world.

### Confidence Judgements

If participants’ confidence was informed by the same prior used to make their perceptual orientation inferences, we hypothesised that their confidence judgements should characterise how well they had matched the low-level statistics in the target with the prior. In contrast, if participants did not use the prior to inform their confidence, their confidence judgements would depend on other heuristic cues, not related to the prior. We tested these predictions with a second model, the confidence model, described below. We first provide an overview of the distribution of confidence responses across response orientations and then outline the assumptions and performance of the confidence model.

The black data in **Figure 3B** shows mean confidence as a function of response orientations. Participants were most confident in their most accurate orientation responses (responses in the 0° bin), suggesting they had some metacognitive awareness of the match of their perceptual judgements to the upright image. There are also clear peaks in confidence at the cardinals (±90° and ±180°), demonstrating that participants’ confidence responses show the same cardinal biases as the perceptual judgements. Our perceptual model revealed that participants’ orientation judgements were consistent with the use of priors for the distribution of orientation energy and phase in natural scenes. We therefore wanted to develop a similar statistical model to determine if participants’ confidence judgements were informed by the same prior.

#### Confidence Model

For the confidence model, we developed several measures that summarised the degree of overlap in orientation energy and phase in the target and the prior. In addition to these prior-related measures, we also considered the possibility that confidence depends on other stimulus cues not related to the prior, such as the contrast of the target patch or the response time of the perceptual judgement (Bertana et al., 2021; Boldt et al., 2017; Faivre et al., 2018; Mole et al., 2018; Patel et al., 2012; Spence et al., 2016; van den Berg, Anandalingam, et al., 2016). To relate these measures to confidence and investigate their relative importance, we used a generalised linear mixed model framework in the form of a modified logistic regression. This allowed us to use a set of weighted stimulus variables to predict binary confidence responses. We describe the assumptions of each predictor in the model and the fixed effect of that predictor below. See **Methods – Confidence Model** for further model description. See **Supplementary Table 1 and Supplementary Figure 2** for model parameters and predictions. See **Supplementary Figure 5** for the correlation between fixed effects.

##### Orientation Energy

We tested two different assumptions about how participants may quantify the degree of overlap between the distribution of orientation energy in the target and the prior. These assumptions were quantified in what we refer to as the *orientation*-*prior-mismatch predictor* and the *cardinal* and *oblique orientation energy predictors*. For the orientation-prior-mismatch predictor, we assumed that participants directly compared a continuous representation of the distribution of orientation features in the stimulus and veridical knowledge of the prior distribution of orientation energy (see **Figure 5A**). The orientation-prior-mismatch predictor, therefore, was a single statistic that summarised the difference between the entire distribution of orientation energy in the target at the rotational offset chosen by the observer and the prior distribution. As shown in **Figure 6B**, however, the effect of the orientation-prior-mismatch predictor on confidence was not significant (*e*^β^ = 0.96, 95% *CI* [0.89, 1.05], *p* = 0.361), suggesting that participants do not appear to explicitly compare the distribution of orientation energy in the target and the prior veridically.

**Figure 5.**
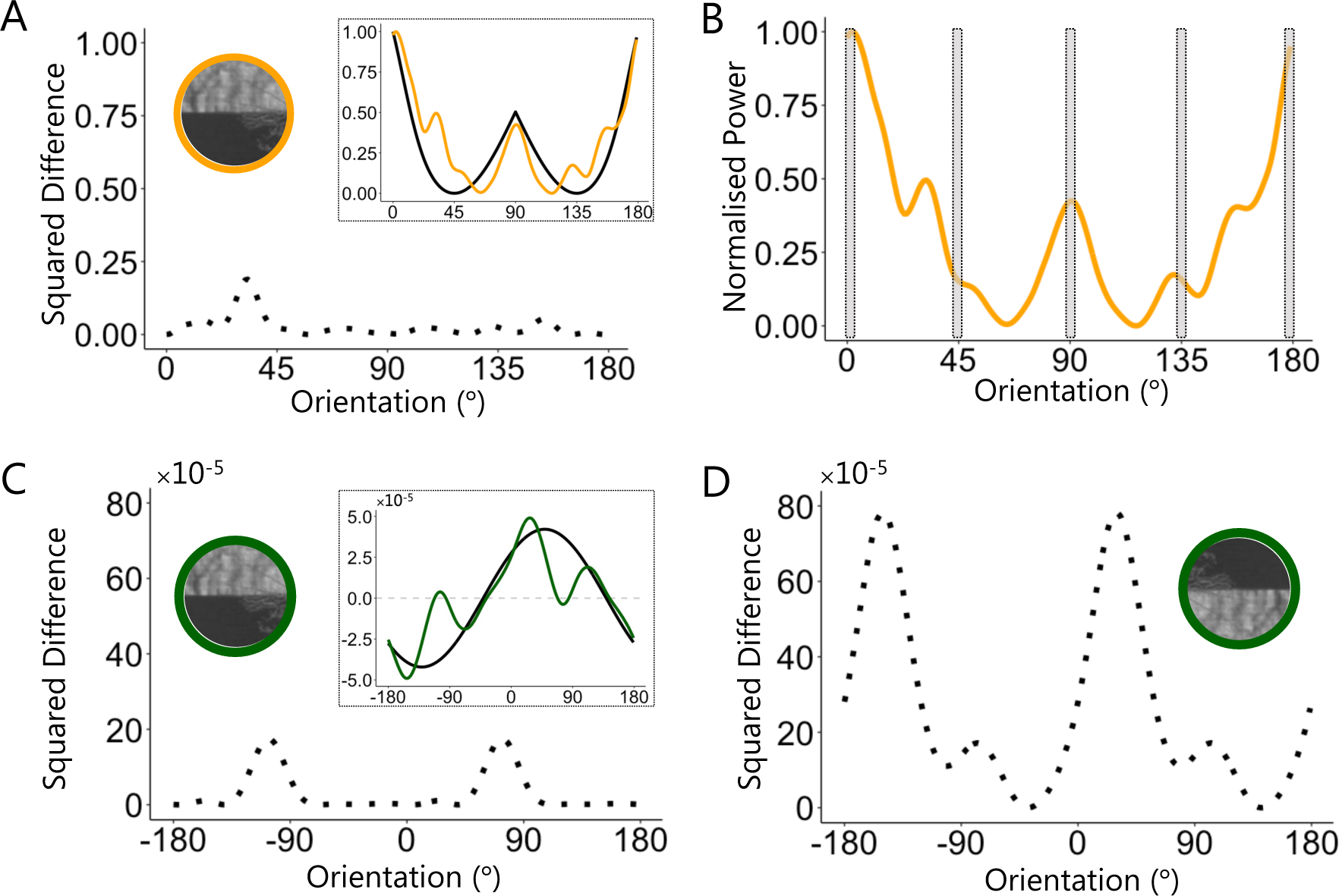
Prior Predictors for Confidence Model. **(A)** Example difference in orientation energy for each bin between the distribution for the target (orange distribution in inset) and the prior (black distribution in inset). The sum of these differences was used as the *orientation*-*prior-mismatch predictor* in the confidence model. **(B)** Example distribution of orientation energy with horizontal (0° or 180°), vertical (90°) and oblique (45° or 135°) energy shaded. **(C)** Squared difference in mean filter response for each filter orientation convolved on the target (green distribution in inset) and across many natural images, the prior (black distribution in inset). The sum of these differences across orientations was used as the *phase-prior-mismatch predictor* in the confidence model. **(D)** As in (C) but for an inverted target. Values were calculated, as in (A) – (D), for each target at the rotational offset chosen by the observer and used to predict confidence.

**Figure 6.**
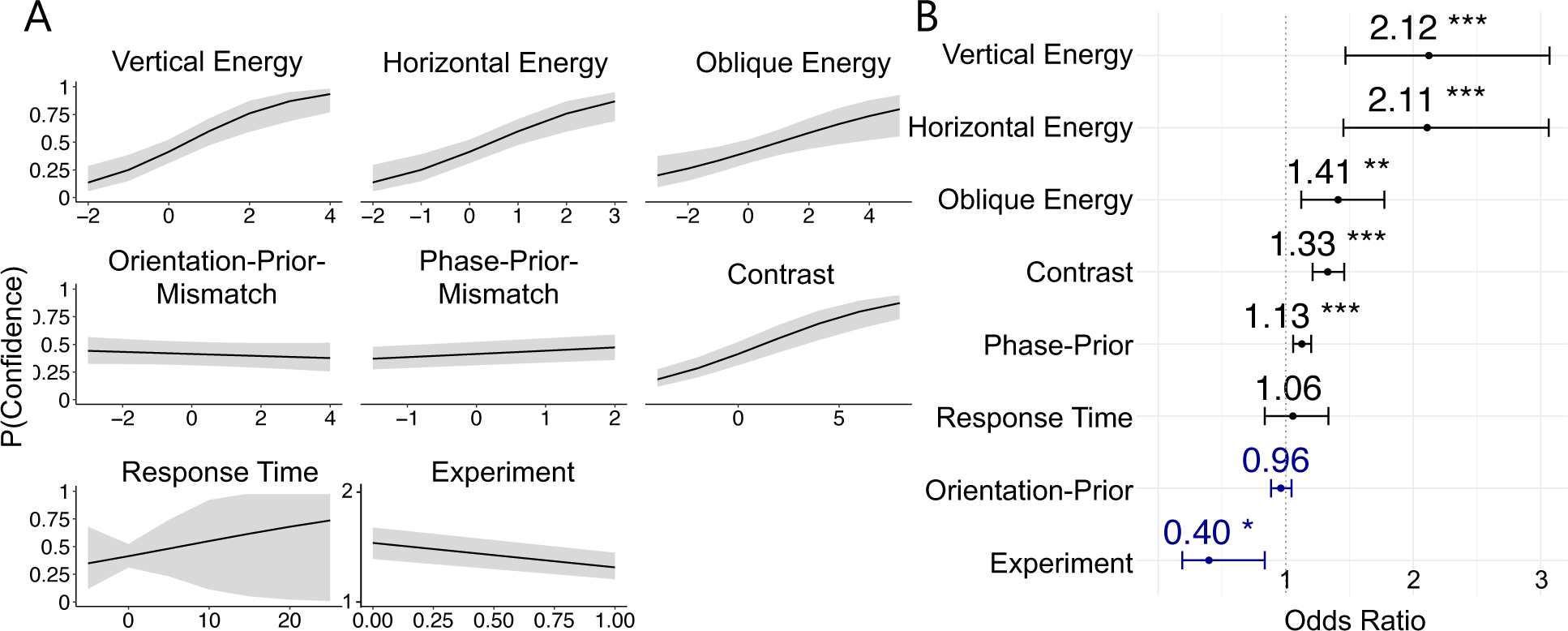
Confidence Model. **(A)** Marginal fixed effects of each predictor on confidence. Shaded regions show ± 1 standard error of the predictions. Note that predictors vertical energy, horizontal energy, oblique energy, orientation-prior-mismatch, phase-prior-mismatch, contrast and response time were standardised. **(B)** Odds ratio for fixed effects. Dotted vertical line indicates odds ratio of 1. Error bars show 95% confidence intervals. Blue values indicate negative coefficients. \*\*\**p* < .001,\*\**p* < .01, \**p* < .05.

As an alternative to computing confidence by comparing the full prior distribution of orientation energy with that measured from the target, we postulated that observers might instead be most confident in targets that contain strong vertical or horizontal features, based on an internalised model of the over-representation of these features in natural scenes. We therefore included a second set of predictors in the confidence model which quantified the amount of vertical (where *θ* = 90°), horizontal energy (where *θ* = 0° *or* 180°) and oblique energy (where *θ* = 45° *or* 135°) in the target at the rotational offset chosen by the participant (see **Figure 5B**). We found that vertical orientation energy (*e*^7^ = 2.12, 95% *CI* [1.47, 3.07], *p* < .001), horizontal orientation energy (*e*^7^ = 2.11, 95% *CI* [1.45, 3.06], *p* < .001) and oblique orientation energy (*e*^7^ = 1.41, 95% *CI* [1.12, 1.77], *p* = .003) were all positively predictive of confidence. The effect of vertical and horizontal energy was almost twice that of oblique orientation energy (see **Figure 6B**), suggesting that participants disproportionally weight the presence of vertical and horizontal orientation features, relative to oblique features, when judging their confidence. This over-weighting is consistent with the distribution of orientation features in natural scenes and suggest that participants confidence is informed by an internalised model of natural image statistics.

##### Phase

Orientation energy is phase invariant, and so a participant cannot use orientation energy alone to distinguish a 0° and ±180° target. We therefore used a predictor in the confidence model that estimated the direction of lighting in the target (see **Methods**). At each filter orientation, we computed the difference between the scaled distribution of phase in the target, as estimated by the phase-locked sine wave filters, and the prior distribution of phase. We squared the difference between the two distributions for each filter orientation and summed across bins so that we had a single measure which summarised the degree of overlap between the phase distribution in the target and the prior. We refer to this measure as the *phase-prior-mismatch* predictor. The effect of the phase-prior-mismatch predictor was significant (*e*^7^ = 1.13, 95% *CI* [1.06, 1.20], *p* < .001). The direction of this effect was positive such that the greater the mismatch between the distribution of phase in the target and the prior, the more confident observers were in their chosen response orientation. This effect appears to be small and we hypothesised that it was driven by participants only using phase information to inform their confidence where there was a strong lighting cue in the target (i.e., where there was a large deviation between the target phase distribution and the prior). Consistent with this hypothesis, in **Supplementary Figure 9** we show that the targets with the greatest deviation from the prior are those with strong lighting cues which do not necessarily originate from the top of the scene. Overall, these findings are consistent with participants relying on an internalised model of statistical regularities in low-level image features in natural scenes, specifically the expected direction of light source, to inform their confidence.

##### Other predictors

We also tested an alternative set of predictions in which participants may rely on learned associations between certain heuristic cues and specific outcomes to judge their confidence. If, contrary to the above effects of orientation energy and phase, participants did not use the prior to inform their confidence, we hypothesised that they might rely on other stimulus features which could approximate task difficulty or sensory uncertainty (Bertana et al., 2021; Boldt et al., 2017; Spence et al., 2016). We therefore included two predictors in the confidence model that were not directly related to the prior: the overall contrast of the target patch (RMS contrast) and the response time of the orientation judgement.

The effect of response time on confidence was not significant (*e*^7^ = 1.06, 95% *CI* [0.84, 1.34], *p* = .721), suggesting that participants do not use response time as a heuristic cue for confidence. The effect of contrast on confidence, however, was significant (*e*^7^ = 1.33, 95% *CI* [1.21, 1.46], *p* < .001), where increasing target contrast was predictive of increasing confidence. Considering the strong effect of oriented contrast energy on confidence reports, it is not particularly surprising that a measure of isotropic orientation was also a significant predictor. Indeed, RMS contrast was correlated with vertical and horizontal energy (see **Supplementary Figure 5**). However, this finding could nonetheless also suggest that, over and above the effect of priors, participants use other heuristic cues to compute their confidence.

To visualise how well these measures accounted for the data more generally, we generated predictions for a model using the statistically significant predictors only (*p* < 0.05 in **Figure 6**; horizontal orientation energy, vertical orientation energy, oblique orientation energy, phase-prior-mismatch and experiment; see **Supplementary Table 2**). The predicted confidence distribution from this model is shown in **Figure 3B** (blue distribution). The model describes the empirical confidence data very well (black distribution; **Figure 3B**), capturing the major peaks in confidence at the cardinal orientations. This finding suggests that the confidence model, with the predictors that quantify the low-level features in the targets and their degree of overlap with the prior and target contrast, provides a compelling recapitulation of how participants judge their confidence in their chosen orientation responses (see **Supplementary Figure 3, Supplementary Table 3, Supplementary Figure 4** for model comparison results).

Despite capturing many aspects of participants’ confidence reports, our model is incomplete. As shown in **Figure 3B**, the confidence model underpredicts confidence for correctly oriented targets (those with response orientations of 0°), suggesting that observers may have had access to other features in the targets, not captured by the model, which led them to confidently infer the correct upright orientation of the target. The model also appears to overpredict confidence for response orientations between ∼65-165°. We expect that this asymmetry in the confidence data, where the pattern of responding is different for response orientations of ∼65-165° and ∼-65-165°, is a result of non-stimulus-specific noise that is not captured by the model. Despite these minor limitations, a small set of environmental statistics provides a reasonable basis from which to understand confidence computations.

## Discussion

We investigated the influence of naturalistic priors on decision confidence. Observers performed a task in which they rotated a small patch of an image of an outdoor scene to its upright orientation and then made confidence judgements in their orientation responses. We found that participants use internal priors about the statistical regularities of low-level visual features in natural scenes to make their perceptual judgements, replicating a recent investigation (A-Izzeddin et al., 2024). Importantly, we also found that participants use the same internal priors to inform their confidence responses, even without explicit instruction about these priors. We discuss the implication of these findings for our understanding of decision confidence below.

### Priors Affect Confidence

We found that participants use prior knowledge about the statistical regularities of orientation energy and phase in natural scenes to inform both their perceptual inferences and confidence judgements.

#### Orientation Energy

Our confidence modelling results showed that the amount of vertical and horizontal orientation energy in the target was an important predictor of confidence. In fact, the effect of vertical and horizontal energy on confidence was almost twice that of oblique orientation energy. Participants were not given any instruction about which image features to use to make their judgements and were not given any explicit instruction about a prior distribution. This finding, therefore, suggests that participants appear to use prior knowledge, either explicit or implicit, about the over-representation of horizontal and vertical orientation features in natural scenes to inform their confidence. This finding broadens our understanding of the influence of the statistical regularities of orientation features in natural scenes on perceptual judgements (Appelle, 1972; Berkley et al., 1975; Campbell et al., 1966; Dakin & Watt, 1997; Girshick et al., 2011; Hansen et al., 2003, 2008; Pratte et al., 2016) and demonstrates, for the first time, that these low-level perceptual priors also effect observers’ confidence in their perceptual decisions.

Although participants appear to use knowledge about the prior probability of certain oriented features in natural scenes to inform their confidence, we did not find evidence that participants directly compare a veridical prior probability distribution of orientation energy in natural scenes and the distribution of orientation energy in the target directly (the *orientation*-*prior-mismatch* predictor) to compute their confidence. Instead, our results suggest that participants use only a subset of orientations to inform their confidence, or, alternatively, use orientations within only some spatial frequencies. This finding suggests that perhaps observers may not use a veridical representation of the prior probability distribution of oriented features but instead, confidence is informed by only the *most predictive* features of the prior and their relative probabilities. As shown in **Figure 7**, perceptually apparent cardinal structures need not be defined by peaks in energy as defined in our prior. When quantifying energy, we aggregated over all spatial frequencies, whereas human vision is known to be bandpass (Campbell & Robson, 1968). Future models could therefore weight an image’s orientation energy according to the human contrast sensitivity. Furthermore, phase alignment at different spatial scales is critical to perceptually relevant edge features that can guide the sorts of perceptual and confidence decisions in our study (Rideaux et al., 2022).

**Figure 7.**
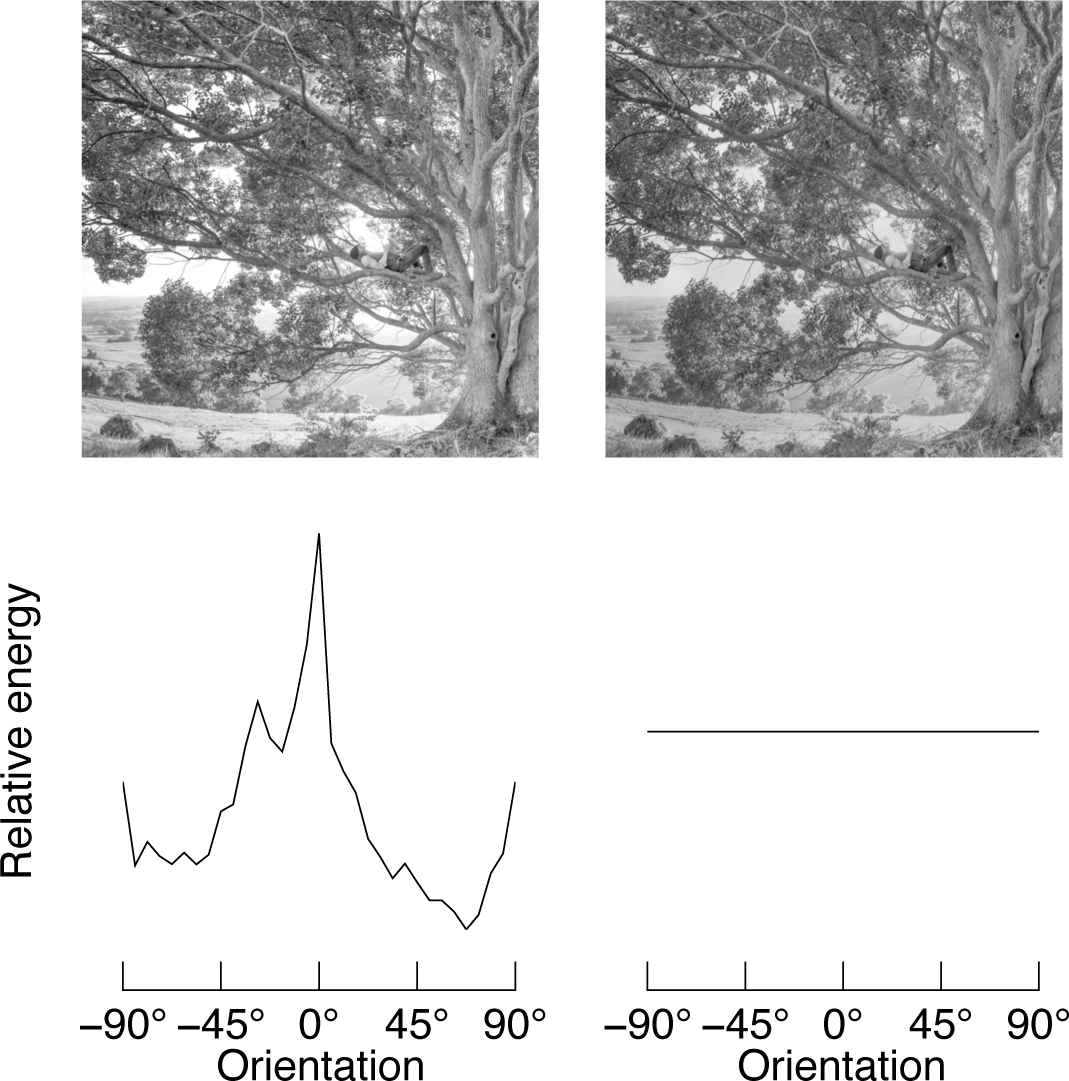
Informativeness of Full Prior Distribution. Comparing the full distribution of orientation energy to a prior may not always be functional. The image on the left shows a greyscale natural image. Its orientation energy is shown below the image, and closely matches the prior distribution shown in Figure 1. The image on the right is the same image, but we have “whitened” its amplitude spectrum with respect to orientation: as shown below the image, the modified image has equal energy at all orientations. Despite the large change in orientation energy, the images are perceptually similar. Because natural images are dominated by low-spatial frequency structures, energy within low spatial frequency bands were over-represented in our measure of orientation energy relative to the bandpass properties of human vision. However, this example nonetheless demonstrates that clearly oriented cardinal structures need not be defined by peaks in energy as defined in our prior per se. Instead, phase alignment at different spatial scales, and orientation energy within different spatial frequency sub-bands, are critical to perceptually relevant edge features that can guide the sorts of perceptual and confidence decisions in our study (Rideaux et al., 2022).

#### Phase

Because orientation energy is phase-invariant, we convolved each target patch with a phase-locked filter to estimate the direction of lighting in the target, consistent with well-documented perceptual effects showing that humans have an internalised model for light originating from above the horizon (Adams, 2007; Brewster, 1826; Metzger, 1936; Murray, 2013; Ramachandran, 1988). We used the difference between the distribution of the phase-locked filter responses in the target at the response orientation chosen by the participant and the distribution across the entire set of images to predict confidence. Specifically, we investigated whether the degree of mismatch between the target phase distribution and the prior, *phase-prior-mismatch*, was predictive of confidence. We found a significant effect of the phase-prior-mismatch predictor. Somewhat unintuitively, however, we observed that the direction of the effect of phase on confidence was such that the larger the mismatch between the prior and target distributions, the more confident participants were in their chosen response orientation. This effect was small. Taken with the results of the perceptual model, we hypothesise that lighting cues are used to orient the targets but that these cues only have a flow on effect to confidence where there are large deviations from the prior. In other words, phase information is most informative for confidence judgements for targets with strong lighting cues that deviate from the expected direction i.e., above. Consistent with this hypothesis, in **Supplementary Figure 9** we show that the targets with the greatest deviation from the prior are those with strong lighting cues which do not necessarily originate from the top of the scene. This effect is again consistent with the fact that participants have internalised models of statistical regularities in the natural world, and can use these models *discriminately* to inform their confidence, i.e., only in cases where lighting is most informative.

To our knowledge, we are the first to show that participants make ambiguous perceptual inferences and judge their confidence in those inferences in a manner that is consistent with comparing a set of image features with an internalised representation of the distribution of those features in the natural world. Our results highlight the role of naturalistic priors for both perception and judgements of confidence and give some insight into the computations underlying the generation of decision confidence.

### The Computational Basis of Confidence

One of the leading theories of confidence suggests that it is computed according to the rules of Bayesian inference where humans combine a *prior* with a *likelihood* to compute a *posterior probability distribution*. Currently, however, the evidence for Bayesian accounts of confidence have been mixed with some studies finding evidence for Bayesian models (Aitchison et al., 2015; Li & Ma, 2020; Navajas et al., 2017; Sanders et al., 2016) and others for non-Bayesian models (Adler & Ma, 2018; Aitchison et al., 2015; Bertana et al., 2021; Denison et al., 2018; Lisi et al., 2021; Locke et al., 2022). One of the major limitations of existing research, however, is that many previous studies have required participants to internalise arbitrary prior distributions (Denison et al., 2018; Li & Ma, 2020; Locke et al., 2022; Qamar et al., 2013; West et al., 2023), making the role of priors in the formation of confidence difficult to interpret. Specifically, if observers are unable to internalise the statistics of a given arbitrary prior distribution within the time-limited context of the experiment, it would be difficult to find evidence of the use of priors in confidence judgements, even if priors are used to inform confidence beyond the lab. In this study, we employed a task design that did not require us to specify a prior distribution. Instead, we relied on participants’ existing priors for natural image statistics that have emerged over evolutionary and developmental timescales (Barlow, 1972; Burgess et al., 1981; Geisler, 2008; Srinivasan et al., 1982).

We found that participants confidence judgements were consistent with an internalised representation of a prior distribution for natural image statistics. While the use of priors is broadly consistent with a Bayesian account of confidence, we did not explicitly test a Bayesian model of confidence in this study. Thus, further model development is required to calculate a full posterior distribution, as per other Bayesian formulations (Aitchison et al., 2015; Denison et al., 2018; Li & Ma, 2020; Locke et al., 2022; Navajas et al., 2017; Sanders et al., 2016; West et al., 2023). Our findings, nonetheless, have important implications for future studies evaluating the computational basis of confidence. Specifically, our results suggest that the use of arbitrary priors in previous research may have obscured the importance of priors in confidence. We caution against using arbitrary prior distributions as participants may be unable or unwilling to internalise the statistics of these distributions, leading to systematic deviations from prior informed behaviour. Overall, our study highlights the importance of naturalistic task designs that capitalise on existing, long-term priors to distinguish among candidate computational models of confidence.

### Other Stimulus Features as Cues for Confidence

In addition to the effect of priors on confidence, we investigated if participants used other cues to inform their confidence. Inconsistent with some previous research, we found that when controlling for other stimulus related cues, response time did not influence confidence (Faivre et al., 2018; Patel et al., 2012; van den Berg, Anandalingam, et al., 2016). Contrary to previous research, we used a task design where responses were continuous (albeit circular) and response execution required incremental adjustment of the response variable. Such a design makes the parsing of decision time and non-decision time (e.g., response execution) challenging. Future research should consider differentiating decision time from non-decision time to better understand the role of response time in the formation of confidence.

We found that the contrast of the target patch had a significant effect on confidence, where higher target contrast was associated with greater confidence. We postulate that stimulus contrast in our experimental paradigm may provide an important cue about sensory uncertainty. Lower contrast targets provide less clear and consistent visual information (Bex & Makous, 2002), and, therefore, in this task contrast can provide a meaningful cue about the perceptual uncertainty of the source of information on which the decision is based. This is consistent with studies showing that confidence is influenced by perceived sensory uncertainty (Adler & Ma, 2018; Denison et al., 2018; Michael et al., 2015; West et al., 2023). Furthermore, it is generally consistent with theoretical work which links confidence with the hypothesis that perceptual uncertainty is encoded as the variance in firing rates across neural populations, with increased uncertainty leading to down-weighting of that evidence source in perceptual integration (Beck et al., 2008; Ma et al., 2006). Further research is required to confirm these hypotheses and understand the neural mechanisms by which certain stimulus features, like image contrast, are involved in the computation of confidence.

### Conclusions

Making adaptive decisions in a noisy perceptual world in the absence of explicit feedback requires not only accurate perceptual judgments, but also accurate judgments of confidence. In this study, we capitalised on the statistical regularities in natural scenes to show that like an optimal observer, participants’ perceptual and metacognitive judgments were tuned to natural image priors. Specifically, our findings support the idea that observers combine multiple features of incoming sensory information, such as sensory uncertainty, with prior probability representations to compute their confidence. Our study highlights the importance of using naturalistic task designs that capitalise on existing, long-term priors to further our understanding of the computational basis of confidence.

## Supporting information

All Supplemental

## Acknowledgements

We thank Cheyanne Gu for assistance with data collection. This work was funded by an ARC DECRA to WJH (DE190100136).

