## Supplementary material for "Priors for natural image statistics inform confidence in perceptual decisions": All Supplemental

### Experiment 1 and Experiment 2 Comparison

Experiment 1 and Experiment 2 were identical with the exception of one minor change in the instruction to participants for how they should report their confidence. In Experiment 1, we asked participants to use high and low confidence responses approximately equally often. We did so because of the unnatural nature of the task – subjectively, it is difficult to infer the upright orientation of the target without any high-level or contextual information of trial-by-trial feedback. We therefore wanted to avoid participants using low confidence responses excessively. That is, we needed variation in confidence responses across targets in order to model the data. In Experiment 2 we wanted to see if we could replicate the same pattern of results without instructing participants about how to use the confidence rating scale. As shown in **Supplementary Figure 1A**, we observed the same patterns in participants' perceptual judgements **(A)** in both experiments. As shown in **Supplementary Figure 1B**, there is a mean decrease in confidence in Experiment 2 (as expected, confidence ratings were lower without an explicit instruction to use high and low ratings equally as often). However, **Supplementary Figure 1C** shows confidence from Experiment 1 and Experiment 2 after removing the mean difference in confidence across experiments, and reveals similarities in the peaks in confidence across response orientation bins. Because the observed pattern of confidence judgements across response orientations was so similar, we combined the datasets and estimated a beta coefficient for an experiment indicator variable in all GLMM confidence models. See **Supplementary Tables 1-3** for the beta coefficients for the experiment indicator variable.

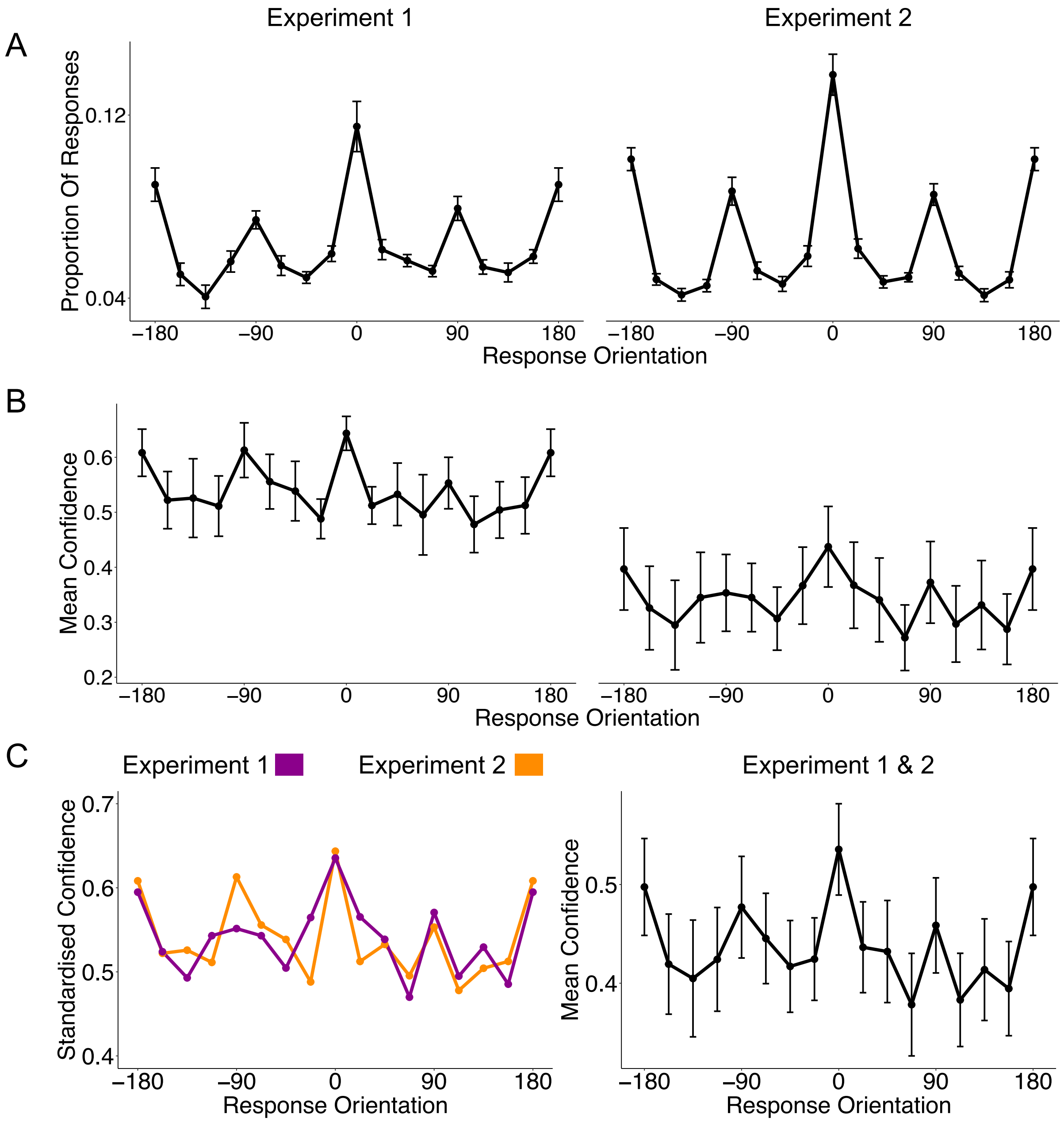


**Supplementary Figure 1. Perceptual and Confidence Judgements Across Experiments.** In Experiment 1, participants were instructed to use high and low confidence responses approximately equally often and in Experiment 2, participants were not given any instructions about how to use the confidence scale. **(A)** Proportion of responses in each response orientation bin for Experiment 1 (left panel) and Experiment 2 (right panel). **(B)** Mean confidence in each response orientation bin for Experiment 1 (left panel) and Experiment 2 (right panel). **(C)** Correcting for mean differences in confidence across studies to visualise similarities in confidence peaks. **(D)** Mean confidence with all data from both experiments. Experimental data as in **Figure 3**. Bin size = 22.5°, $N_{Experiment1}$ = 10, $N_{Experiment2}$= 11, Error bars show ± 1 *SEM*.

**Supplementary Table 1**

***GLMM Parameters and Model Statistics for Full Confidence Model***

| Predictors | Odds Ratios | *95% Confidence Interval* | *p** |
|---|---|---|---|
| (Intercept) | 1.14 | 0.63 – 2.06 | 0.659 |
| Vertical Orientation Energy | 2.12 | 1.47 – 3.07 | **<0.001** |
| Horizontal Orientation Energy | 2.11 | 1.45 – 3.06 | **<0.001** |
| Oblique Orientation Energy | 1.41 | 1.12 – 1.77 | **0.003** |
| Orientation Prior Mismatch | 0.96 | 0.89 – 1.05 | 0.361 |
| Phase Prior Mismatch | 1.13 | 1.06 – 1.20 | **<0.001** |
| Contrast | 1.33 | 1.21 – 1.46 | **<0.001** |
| Response Time | 1.06 | 0.84 – 1.34 | 0.646 |
| Experiment | 0.40 | 0.19 – 0.84 | **0.015** |
| **Overall Model Statistics** | | | |
| Marginal $R^2$ / Conditional $R^2$ | 0.079 / 0.363 | | |
| *AIC* | 11993.20 | | |
| *BIC* | 12319.86 | | |

* Significant p-values are bolded

**Model Predictions for the Full Confidence Model**

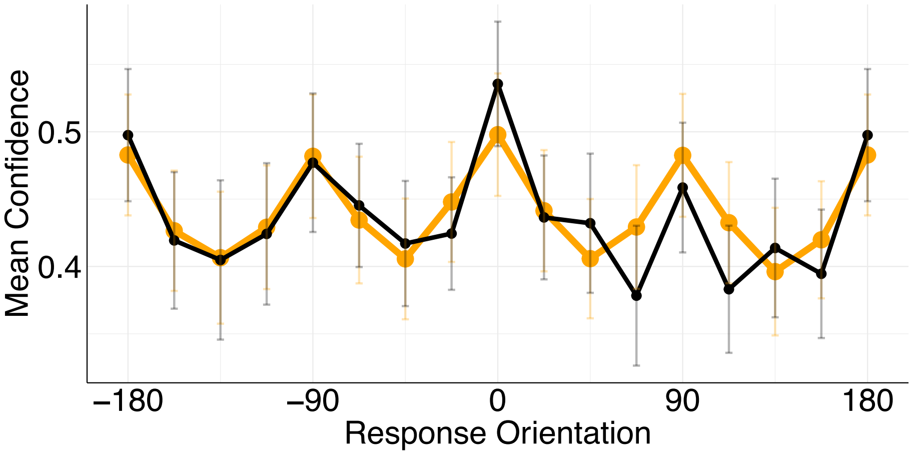


**Supplementary Figure 2. Full Confidence Model.** The proportion of high confidence responses across binned response orientations is shown in black. The output of the model with all predictors (See **Supplementary Table 1**) is shown in orange. Bin size = 22.5°. Error bars show ± 1 *SEM*.

### Supplementary Table 2

**GLMM Parameters and Model Statistics for Confidence Model with Significant Predictors Only**

| Predictors | Odds Ratios | *95% Confidence Interval* | *p** |
|---|---|---|---|
| (Intercept) | 1.17 | 0.66 – 2.07 | 0.588 |
| Vertical Orientation Energy | 2.40 | 1.60 – 3.59 | **<0.001** |
| Horizontal Orientation Energy | 2.44 | 1.61 – 3.69 | **<0.001** |
| Oblique Orientation Energy | 1.49 | 1.18 – 1.88 | **0.001** |
| Phase Prior Mismatch | 1.12 | 1.06 – 1.19 | **<0.001** |
| Contrast | 1.33 | 1.22 – 1.45 | **<0.001** |
| Experiment | 0.36 | 0.18 – 0.74 | **0.005** |
| **Overall Model Statistics** | | | |
| Marginal $R^2$ | 0.121 | | |
| *AIC* | 12149.37 | | |
| *BIC* | 12352.621 | | |

* Significant p-values are bolded

### Supplementary Table 3

**GLMM Parameters and Model Statistics for Intercept Model**

| Predictors | Odds Ratios | *95% Confidence Interval* | *p** |
|---|---|---|---|
| (Intercept) | 1.26 | 0.71 – 2.24 | 0.425 |
| Experiment | 0.35 | 0.16 – 0.78 | **0.010** |
| **Overall Model Statistics** | | | |
| Marginal $R^2$ / Conditional $R^2$ | 0.061 / 0.253 | | |
| *AIC* | 12657.19 | | |
| *BIC* | 12678.97 | | |

* Significant p-values are bolded

### Model Predictions for Intercept Only Model

To provide a reference point for model performance, we fit an intercept only model with an intercept term for each participant (see **Supplementary Figure 3** and **Supplementary Table 3**). We found substantially poorer performance for the intercept model ($AIC_{intercept}$ = 12657.19, $BIC_{intercept}$ = 12678.97, $AIC_{conf}$ = 12149.46, $BIC_{conf}$ = 12352.71; see **Supplementary Figure 4**).

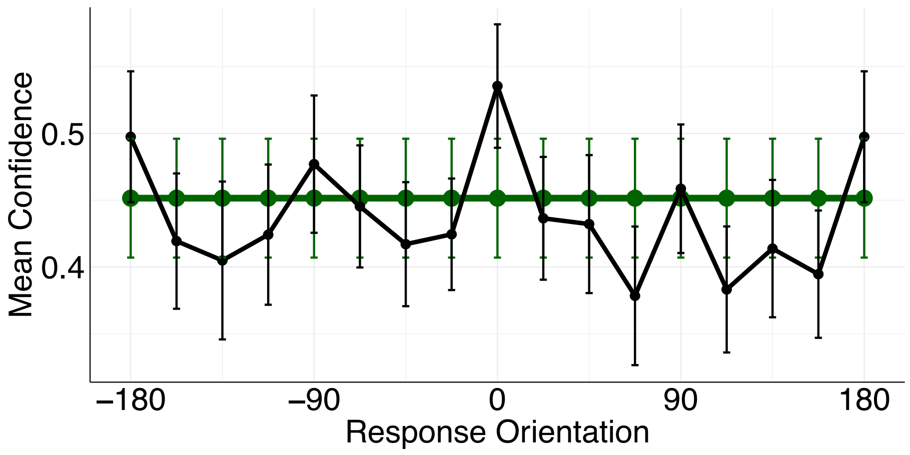


**Supplementary Figure 3. Intercept Only Model.** The proportion of high confidence responses across binned response orientations is shown in black. The output of the intercept model (See **Supplementary Table 3**) is shown in green. Bin size = 22.5°. Error bars show ± 1 *SEM*.

**Model Comparison**

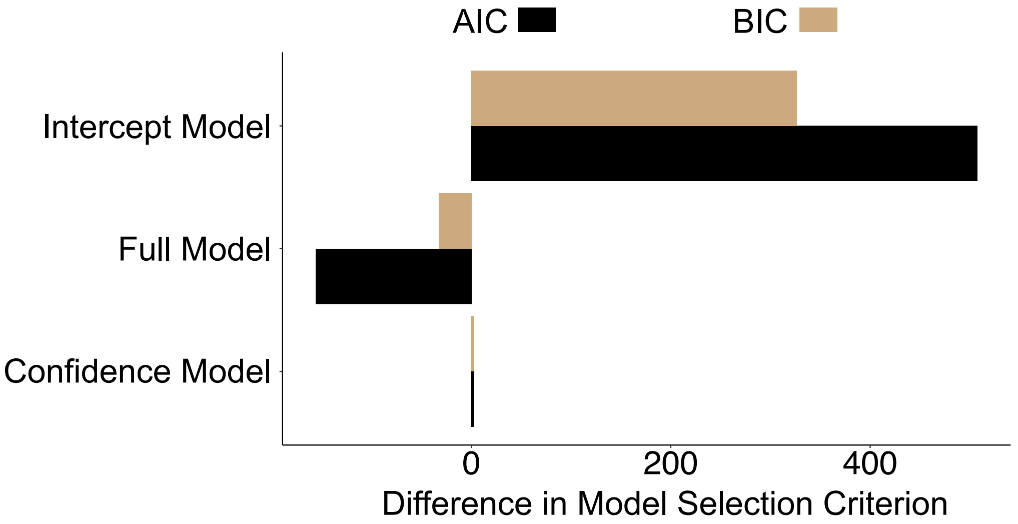


**Supplementary Figure 4. Model Comparison.** Differences in *AIC* and BIC from main confidence model (model with significant predictors only). Negative values indicate preference for model on y axis. The full model (with all predictors) had the lowest *AIC* and *BIC* score.

**Supplementary Table 4**
**Noise Parameters from the Perceptual Model**

| Subject | Sigma Parameter |
|---|---|
| 1 | 19.68 |
| 2 | 22.75 |
| 3 | 21.54 |
| 4 | 21.16 |
| 5 | 24.35 |
| 6 | 37.88 |
| 7 | 25.42 |
| 8 | 29.14 |
| 9 | 37.23 |
| 10 | 35.78 |
| 11 | 21.70 |
| 12 | 21.71 |
| 13 | 27.90 |
| 14 | 18.97 |
| 15 | 22.80 |
| 16 | 22.15 |
| 17 | 22.93 |
| 18 | 19.43 |
| 19 | 21.19 |
| 20 | 25.28 |
| 21 | 23.52 |

### Correlation of Fixed Effects

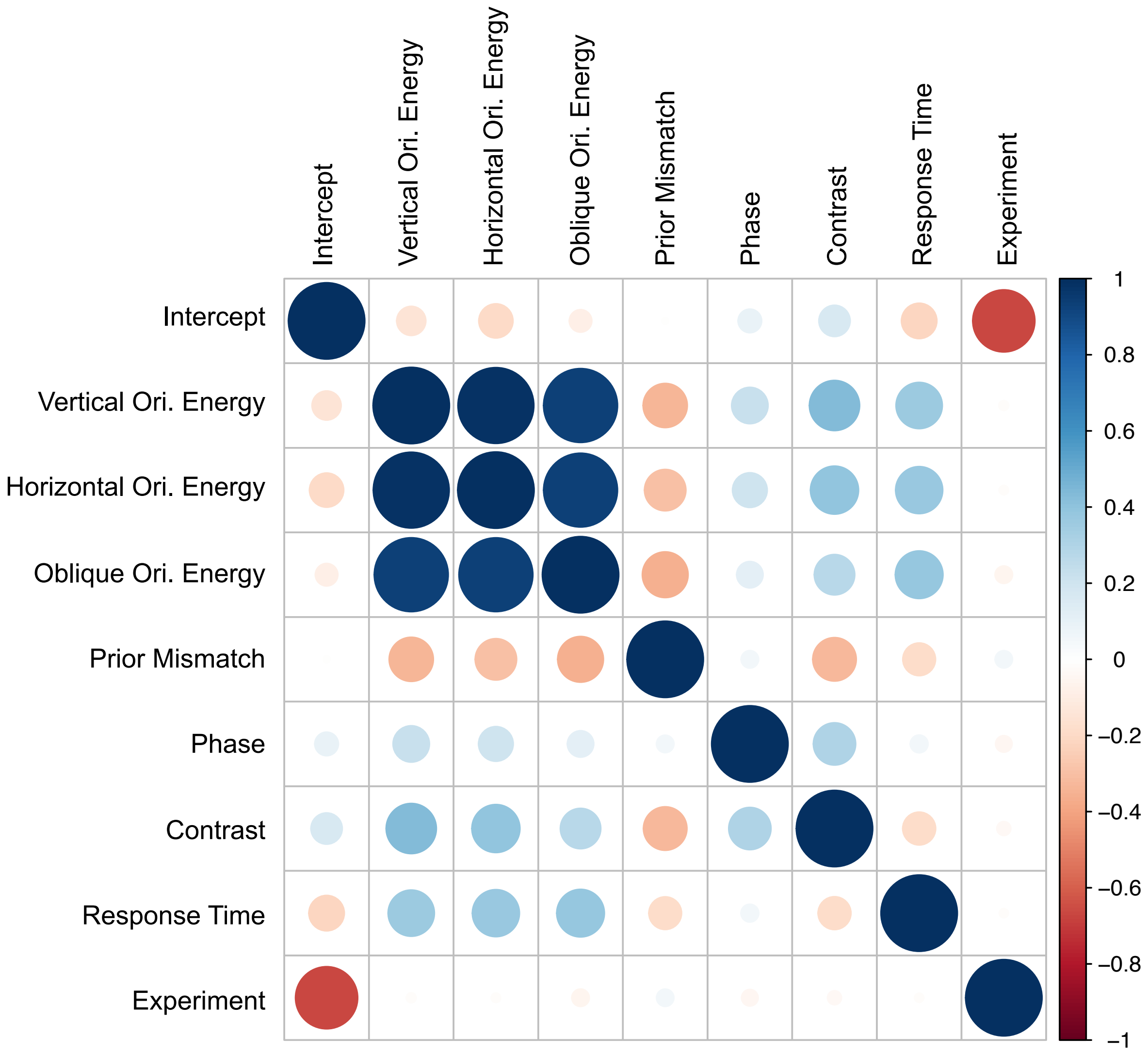


**Supplementary Figure 5. Correlations for Fixed Effects.** Correlation matrix for the fixed coefficient estimates in the confidence model. There was a high correlation between the fixed effects for the orientation energy predictors. This was expected given the divisive-normalisation-like procedure (Carandini & Heeger, 2012). See **Supplementary Figure 6**.

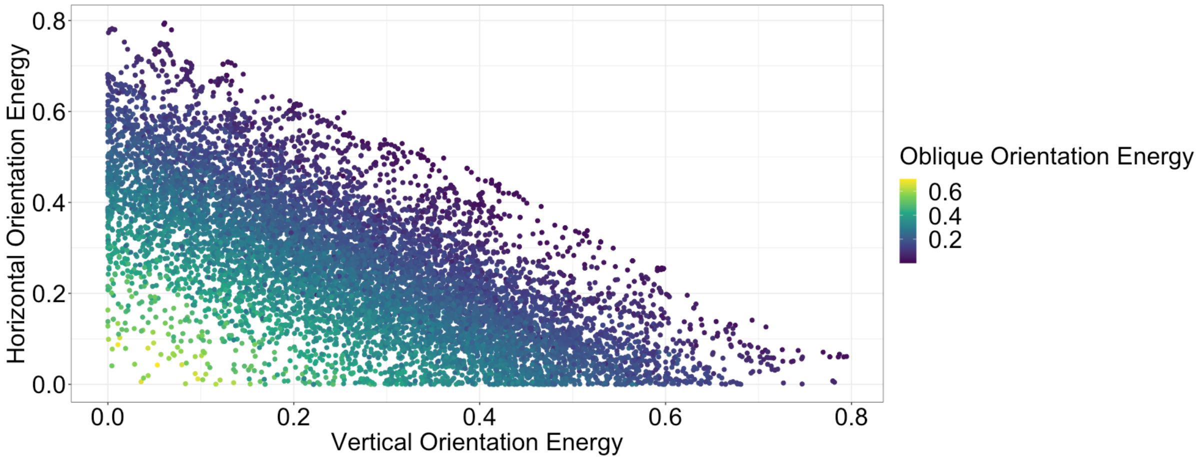


**Supplementary Figure 6. Relationship Between Orientation Energy Predictors.** We used a divisive-normalisation-like procedure for the orientation energy predictors in the confidence model. This meant that each orientation band expressed a proportion of orientation energy in that bin relative to total orientation energy in the target.

### Frequency Distributions for Confidence Predictors

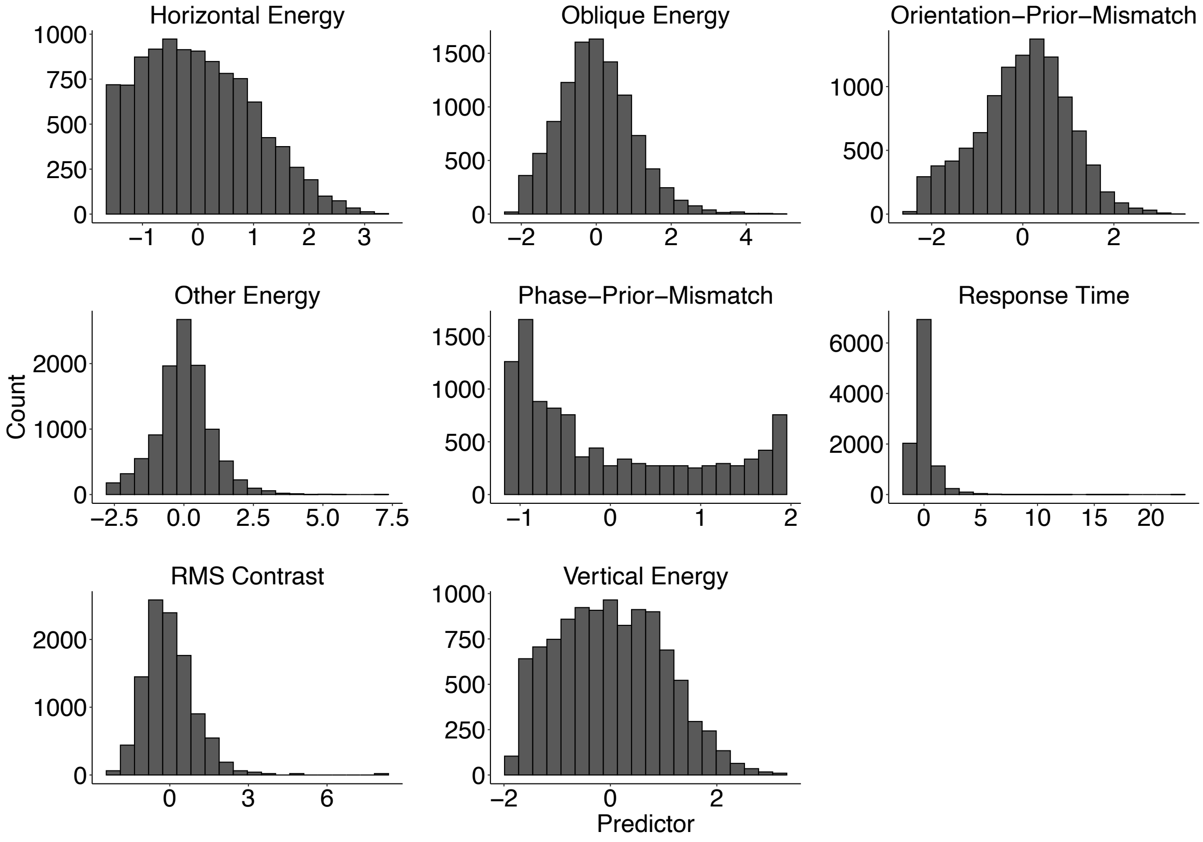


**Supplementary Figure 7. Frequency Distributions for Confidence Predictors.** Frequency distributions for standardised predictors in the confidence model. Predictors were standardised by subtracting the mean and dividing by the standard deviation of all values.

### High-Level Image Features and “Informativeness” Data

Targets were selected randomly from a database and were not screened before being shown to participants. Thus, the possibility remained that a subset of targets may have contained high-level image features (e.g., letters, objects) which could be used to unambiguously cue the objective upright orientation of the targets. We therefore wanted to confirm that the observed data and modelling results were not driven by responses to targets that contained high-level features. We investigated this question using informativeness ratings, described below, from a different experiment which used the same targets (A-Izzeddin et al., 2024).

#### *Informativeness Study*

We used informativeness ratings from A-Izzeddin et al. (2024). See **Control Experiment 1** in **Supplementary Materials** of A-Izzeddin et al. (2024) for full experimental details. We provide a brief overview of the experimental protocol below to give sufficient context for our analyses.

**Methods.** On each trial, participants (*N* = 2; Authors RKW and EJA) were shown a target in its upright position and were instructed to categorise the target as “informative” or “uninformative”. Specifically, participants were instructed to judge whether there was sufficient high-level information in the target to unambiguously indicate the “correct” upright orientation of the target. For example, patches that contained identifiable objects such as cars or signs were to be classified as informative. One participant was instructed to use a liberal response criterion (P1 rated 295 out of 7176 targets as informative) and the other was instructed to use a conservative response criterion (P2 rated 66 out of 7176 targets as informative). Participants categorised a random subset of all possible targets from a larger database (including targets not tested in this study), which meant that we had informativeness data for 370 out of the total 500 targets shown to participants in Experiment 1 and 2. We categorised a target as informative if either (or both) participants rated that target as being informative and used these categorisations to further investigate the confidence data from Experiment 1 and Experiment 2.

**Results.** Of the 500 targets shown to participants in Experiment 1 and Experiment 2: 71.4% were categorised as *not* containing sufficient high-level features to unambiguously inform the upright orientation of the target; 2.6% were categorised as having informative high-level features; and the remaining 26% were not categorised.

We show the confidence data for targets with informativeness ratings in **Supplementary Figure 8.** Importantly, as shown in **Supplementary Figure 8A**, for targets rated as *not informative*, we still see the characteristic peaks in confidence at the cardinal orientations in the empirical data. This finding suggests that the observed patterns in the empirical data shown in **Figure 3B**, were not driven by responses to targets containing high-level features. Furthermore, there remains a close correspondence between the predictions of the confidence model and the empirical data. For targets rated as *informative*, as shown in **Supplementary Figure 8B**, there was less overlap between model predictions and the empirical data, suggesting that when the targets contain sufficient high-level features, participants relied less on priors for low-level features to compute their confidence.

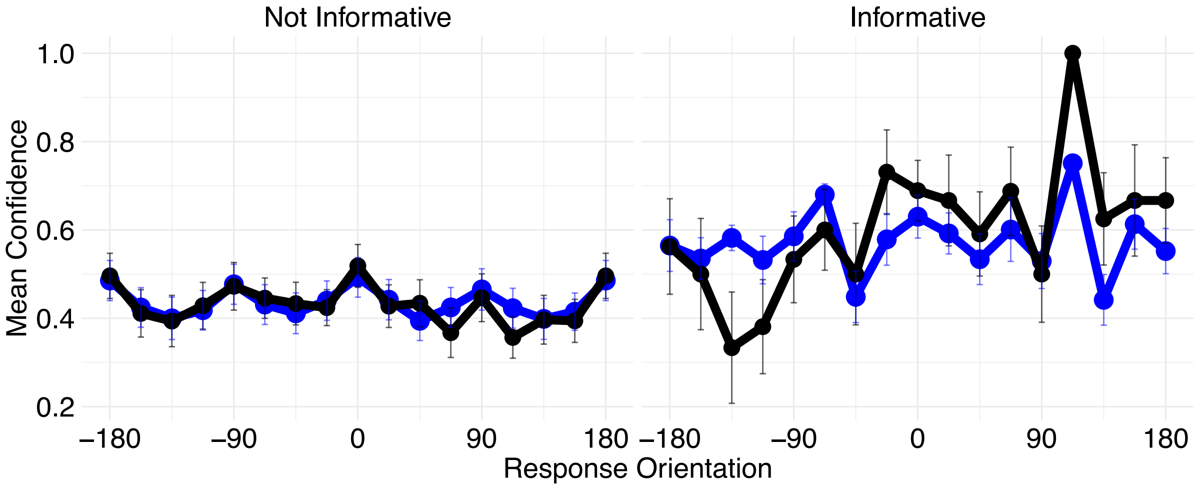


**Supplementary Figure 8. High-Level Image Features and Informativeness Ratings.** Mean confidence across binned response orientations is shown in black and the output of the confidence model is shown in blue for targets rated as *not informative* (left) and *informative* (right).

Taken together, these results indicate that participants may have had access to sufficient high-level features which they used to judge their confidence in a very small proportion of images. However, there was neither sufficient numbers of informative images shown to participants, nor consistent pattern of responses to informative images to account for the observed data.

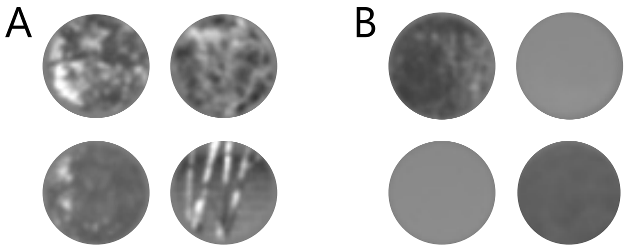


**Supplementary Figure 9. Phase-Prior-Mismatch Target Comparison. (A)** Targets with the greatest difference between the measured distribution of phase and the prior distribution. Differences were calculated using the difference between a set of phase-locked filters at orientations 0 – 179° convolved on the target and compared with the average distribution across the entire image set. **(B)** Targets with the smallest difference between the measured distribution of phase and the prior. These examples make clear that our phase estimate is not always a predictive cue of lighting direction.

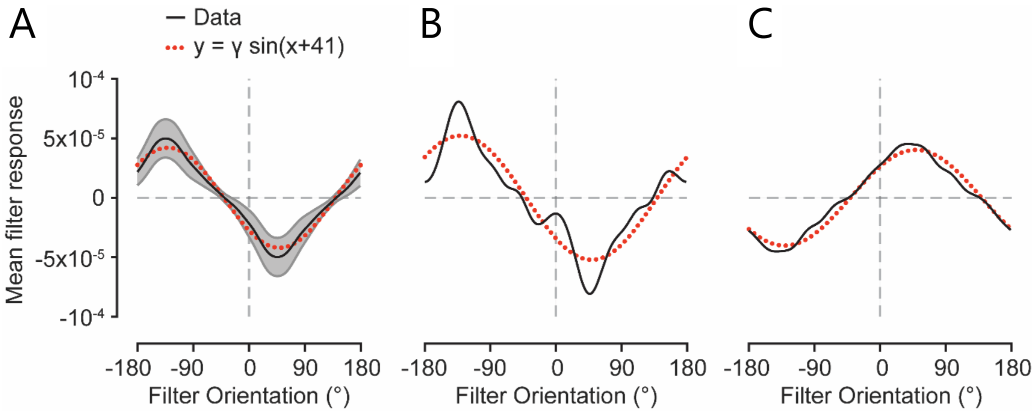


**Supplementary Figure 10. Estimating Lighting Direction via Phase**. **(A)** Phase-dependent contrast as a function of filter orientation, averaged over 9360 target patches. We convolved each target patch with a sine wave filter, and then found the average of the filter response within the centre of the target patch to exclude edge artifacts. The mean filter response and 95% confidence intervals across all target patches are shown by the black line and shaded region, respectively. The dotted red line is the best fitting sinusoidal function, which captures 97% of the variance of the mean filter response. The sinusoid is offset by 41°; the value, $\gamma$, is a scaling factor. **(B-C)** To model observers' final orientation report, we regressed the best fitting sinusoidal function, shown in (A), onto the phase distribution for each target patch. Note that, if the sign of the scaling factor is negative as it is in (C), then the target patch needs inversion for it to best match the aggregate phase distribution. Units in (B) and (C) are arbitrary. Original figure and caption from A-Izzeddin et al. (2024).
